# No metabolic “default mode” of human brain function

**DOI:** 10.1101/2020.08.17.245183

**Authors:** Lars S Jonasson, Filip Grill, Andreas Hahn, Lucas Rischka, Rupert Lanzenberger, Vania Panes Lundmark, Katrine Riklund, Jan Axelsson, Anna Rieckmann

## Abstract

The finding of reduced activity in the default mode network (DMN) during externally focused cognitive control has been highly influential to our understanding of human brain function, but ‘deactivations’ have also prompted major questions of interpretation. Using hybrid functional PET-MR imaging, this study shows that fMRI task activations and deactivations do not reflect antagonistic patterns of synaptic metabolism. FMRI activations were accompanied by concomitant increases in metabolism during cognitive control, but, unlike the BOLD response, metabolism in the core DMN did not change between rest and task. Metabolic increases along the borders of the DMN during task performance further revealed a set of regions that guide engagement and suppression of neighboring networks during cognitive control. Collectively, dissociations between metabolism and BOLD signal specific to the DMN reveal functional heterogeneity in this network and demonstrate that BOLD deactivations during cognitive control should not be interpreted to reflect reduced synaptic activity.

## Introduction

More than 20 years ago, Shulman and colleagues described a set of brain regions that consistently showed low levels of functional brain activity during cognitive control tasks relative to a passive resting state^1^. In their seminal paper, they reported data from nine [^15^O]-*positron emission tomography* (PET) experiments with passive or low-level baseline conditions as well as active task conditions including either language or non-language cognitive tasks. By subtracting baseline scans from active task scans they were able to identify areas displaying decreased blood-flow during the active task conditions, later conceptualized as a resting or “default” state of brain function^2,3^.

Since its early description as task-related deactivations in PET-based measures of blood flow and oxygen consumption, activity in the *default mode network* (DMN) has been explored extensively using *functional magnetic resonance imaging* (fMRI) and its function remains a topic of great interest in the cognitive neurosciences^4–6^. fMRI work showed that an antagonistic relationship between the task-negative DMN and task-positive association networks, including the *dorsal attention network* (DAN), is also evident in functional connectivity studies without task, suggesting that it is an intrinsic feature of functional brain network architecture^7,8^.

A dominant view posits that task deactivations in DMN reflect a relative increase of activity in the DMN during rest, related to its role in unconstrained, internally directed thought not driven by sensory input, including autobiographical thoughts, thinking about past and future, and mind wandering^9–13^. However, recent work has begun to emphasize a more general role for the DMN in ongoing regulation and reconfiguration of brain dynamics and network states in varying contexts^4,14–17^. On this view, decreased activity during task is reflective of such reconfigurations being more prominent during an unconstrained state. There is also substantial evidence that *blood-oxygenation-level-dependent* (BOLD) deactivations in the DMN during external cognitive control are relevant to understanding human cognition in disease. Individual differences in the strength of negative coupling between networks during cognitive control have been linked to task performance^18^, and a failure to deactivate the DMN has been associated with increased attentional lapses^19^, reduced task-performance in aging^20–22^, Alzheimer’s disease^21,23^, and schizophrenia^24,25^.

Despite the large influence that the DMN deactivations described by Shulman and colleagues have had on subsequent research and our understanding of large-scale association networks in human cognition, the division of large-scale cortical association networks into antagonistic task-positive and task-negative networks has almost exclusively been demonstrated with non-invasive imaging modalities that depend on neurovascular coupling (i.e. fMRI, [^15^O]-PET and arterial spin labeling), leaving it uncertain whether deactivations and activations during externally focused cognitive control tasks reflect antagonistic patterns of synaptic activity.

The current understanding of the neurovascular coupling underlying negative BOLD responses during task and whether they follow the same principles as positive BOLD signals is limited. Invasive recordings of neural activity in animals combined with BOLD imaging have studied the negative BOLD response primarily in early sensory-motor regions and hippocampus with conflicting results^26–30^. The limited number of animal studies on DMN deactivations have also failed to show clear evidence that BOLD task deactivations in DMN are reflective of relative decreases in neural activity during externally focused cognitive control^31–35^.

[^18^F]*fluorodeoxyglucose* (FDG) PET offers a non-invasive imaging technique with whole brain coverage at millimeter resolution that provides a measure of synaptic activity that is independent of neurovascular coupling. In resting-state FDG scans, the DMN can be identified as a network with high metabolic activity, particularly in the *posterior cingulate cortex* (PCC) and precuneus^2,36,37^. Fox et al. first showed that FDG is also sensitive to local task demands across scans^38^. Recently, FDG imaging has been advanced to show that transient changes in glucose metabolism can be measured dynamically in a single scan in human subjects^39,40^. This *functional PET* (fPET) method relies on a modification of the traditional bolus injection paradigm into a slow infusion protocol that delivers a constant plasma supply of FDG to the cell, such that a dynamic change in the slope of the time activity curve is proportional to the rate of CMR_glu_. Using this method, transient metabolic changes over task blocks that are only a few minutes long have been detected in task-relevant regions during visual stimulation, hand movement as well as visuo-spatial reasoning^39–44^, demonstrating marked spatial overlap with BOLD responses during the same task^44^.

In the current study, we use simultaneous fPET-fMRI to study concurrent changes in blood oxygenation and glucose metabolism during seven alternating blocks of rest and a *working memory* (WM) task. This task elicits robust BOLD activations in core WM regions within the *fronto-parietal control network* (FPN) and DAN, and deactivations in the DMN (Fig. 1 and Online Methods). The primary aim is to test the hypothesis that task-positive and task-negative BOLD changes during this externally focused cognitive control task are physiological opposites also in terms of glucose metabolism. The results will inform us whether BOLD deactivations in DMN during externally focused cognitive control compared to rest should be interpreted as reduced neural activity.

**Fig. 1.**
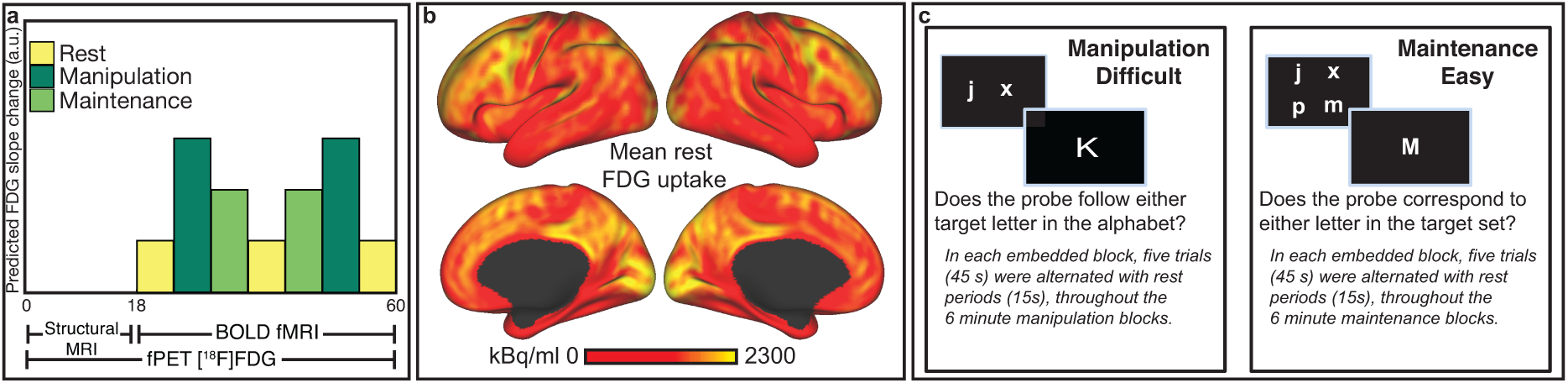
Experimental setup. **a** Constant infusion of FDG started at time 0 and continued for 60 min. The FDG uptake was modelled using the *general linear model* (GLM) with separate regressors for the manipulation (difficult) and maintenance (easy) working memory tasks in the shape of ramp functions that increase by one during the respective 6-minute task blocks (i.e. after the two manipulation blocks that regressor has ramped up to 12). Structural images were collected during the first 18 min, and fMRI-BOLD data were collected between 18 and 60 min. To provide suitable contrast for both fPET and BOLD, an embedded block design is used^1^, where each “slow” 6 min task block has six “fast” 1 min blocks embedded within, each involving 45 s task and 15 s rest. **b** Mean uptake of FDG averaged across subjects over the first 24 min task-free period. **c** The manipulation and maintenance working memory tasks.

## Results

BOLD-fMRI and FDG-fPET data were acquired from 23 healthy young adults while they alternated between resting with their eyes open and performing two conditions of a WM tasks (an overview of the experimental setup is shown in Fig. 1). Participants performed both tasks with high accuracy (WM manipulation, M = 92.97%, SD = 5.22; WM maintenance, M= 97.25%, SD = 3.00) but WM manipulation was more difficult compared to WM maintenance (*t*(22) = 4.02, *p* < 0.001). The main results are based on conventional *general linear model* (GLM) analyses contrasting the difficult WM manipulation condition to rest and to the easier WM maintenance condition. To aid the interpretation of the main results, additional analyses include kinetic modelling of fPET, BOLD functional connectivity, and *regions-of-interest* (ROI)-based BOLD time-series analyses.

### fMRI activations and increases in glucose metabolism overlap in the frontoparietal control and attention networks during working memory

We first tested the hypothesis that WM-induced BOLD and FDG activations would show a high degree of overlap in the core task networks. Fig. 2a shows the patterns for fMRI and fPET activations, as well as their spatial overlap Fig. 2b. Both modalities showed significant increases comparing manipulation > rest in canonical WM regions belonging to the FPN, including *dorsolateral PFC* (DLPFC), *ventrolateral PFC* (VLPFC), *anterior cingulate cortex* (ACC), and *superior parietal lobule* (SPL) as well as the DAN, including precentral gyrus and intraparietal sulcus. See Supplementary Data Table 1 for a complete list of peak activation loci for fMRI and fPET. Importantly, both modalities also demonstrated an increase in activity with an increase in task demand (manipulation > maintenance) across the task-positive networks (Fig. 2a, bottom row), in line with recent results from a visuo-spatial task^44^. Across contrasts, this shows that changes in glucose metabolism parallel BOLD signal increases in task-positive regions, providing important initial support for the feasibility of the method and current design.

**Fig. 2.**
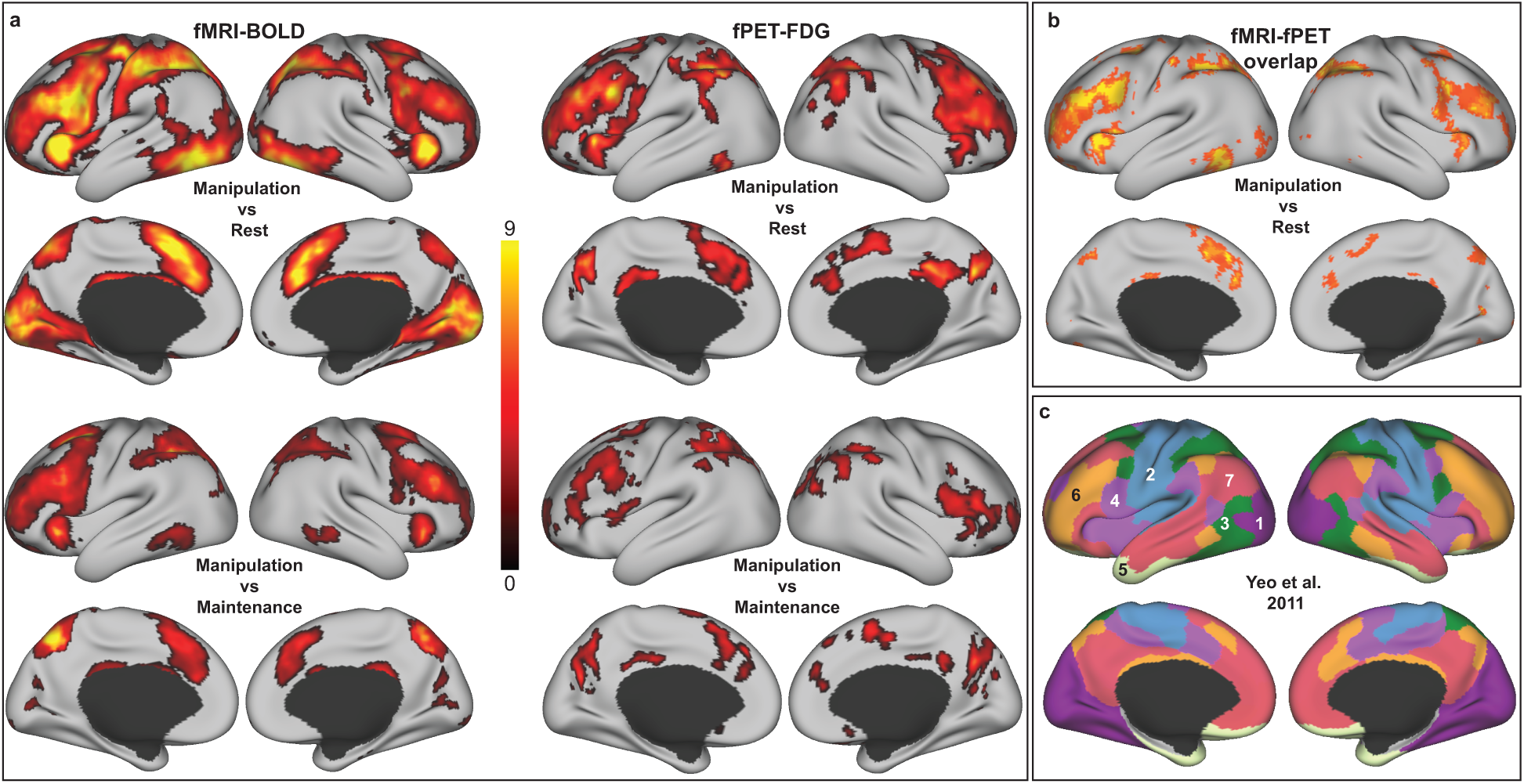
Task-related increases in BOLD signal and glucose metabolism. **a** *t*-maps (corrected at *p* < 0.05) for fMRI (left) and fPET (right) were overlaid on an inflated surface in workbench view. Separate maps are shown for manipulation (the difficult working memory condition) compared to rest (top), and for the contrast manipulation > maintenance (bottom). Both modalities showed significantly larger task-induced responses to manipulation compared to maintenance. **b** Overlap between fMRI and fPET at different uncorrected *t*-thresholds for the manipulation task (p < 0.01 to p < 0.0001, red to yellow, respectively) are modest in visual and sensorimotor networks at lower thresholds but high convergence in FPN and DAN also at more stringent thresholds. **c** The Yeo atlas is displayed to provide a point of reference for the locations of functional activations (1 = visual; 2 = sensorimotor; 3 = dorsal attention; 4 = ventral attention; 5 = limbic; 6 = frontoparietal; 7 = default mode network).

Interestingly, although largely overlapping activations for both modalities were observed in the FPN for manipulation > rest (Fig. 2b with reference parcellation in Fig 2c), large BOLD signal changes in visual, motor, and sensorimotor cortices due to the sensory-motor demands of the task were not accompanied by FDG changes to the same extent; for FDG, task-related increases in visual cortex and left motor cortex were positive but non-significant at the a-priori selected, corrected, threshold. The overlap map in Fig. 2b therefore shows the spatial overlap at different uncorrected thresholds, *p* < 0.01 to *p* < 0. 0001. In order to quantify the partial overlap between different functional networks and provides additional statistical support that voxels with FDG activations overlapped with those showing increased BOLD primarily in FPN, DAN, and VAN, but also that FDG activations are more focal than the widespread task activations in BOLD, overlap was computed separately for activations within a priori defined functional divisions in Supplementary Fig. 1 (see also Supplementary Data Table 1).

### Task-related fMRI deactivations in the DMN are not accompanied by overlapping decreases in glucose metabolism

The next set of analyses explored whether a relative decrease in fMRI activity during task as compared to rest would also be paralleled by task-dependent decreases in FDG uptake. Because both modalities demonstrated the largest change in activation for the contrast manipulation (difficult) > rest, the analyses in the following sections are based on the negative signal changes in this contrast, shown in blue in Fig. 3a and 3b side by side for each modality. In fMRI, the DMN showed widespread deactivation during manipulation as compared to rest, including PCC, *medial prefrontal cortex* (MPFC), inferior lateral parietal cortex, lateral temporal cortex. Strikingly, aside from temporal cortex (explored further in Supplementary Fig. 2), no spatially overlapping changes in glucose metabolism were observed in core regions of the DMN.

**Fig. 3.**
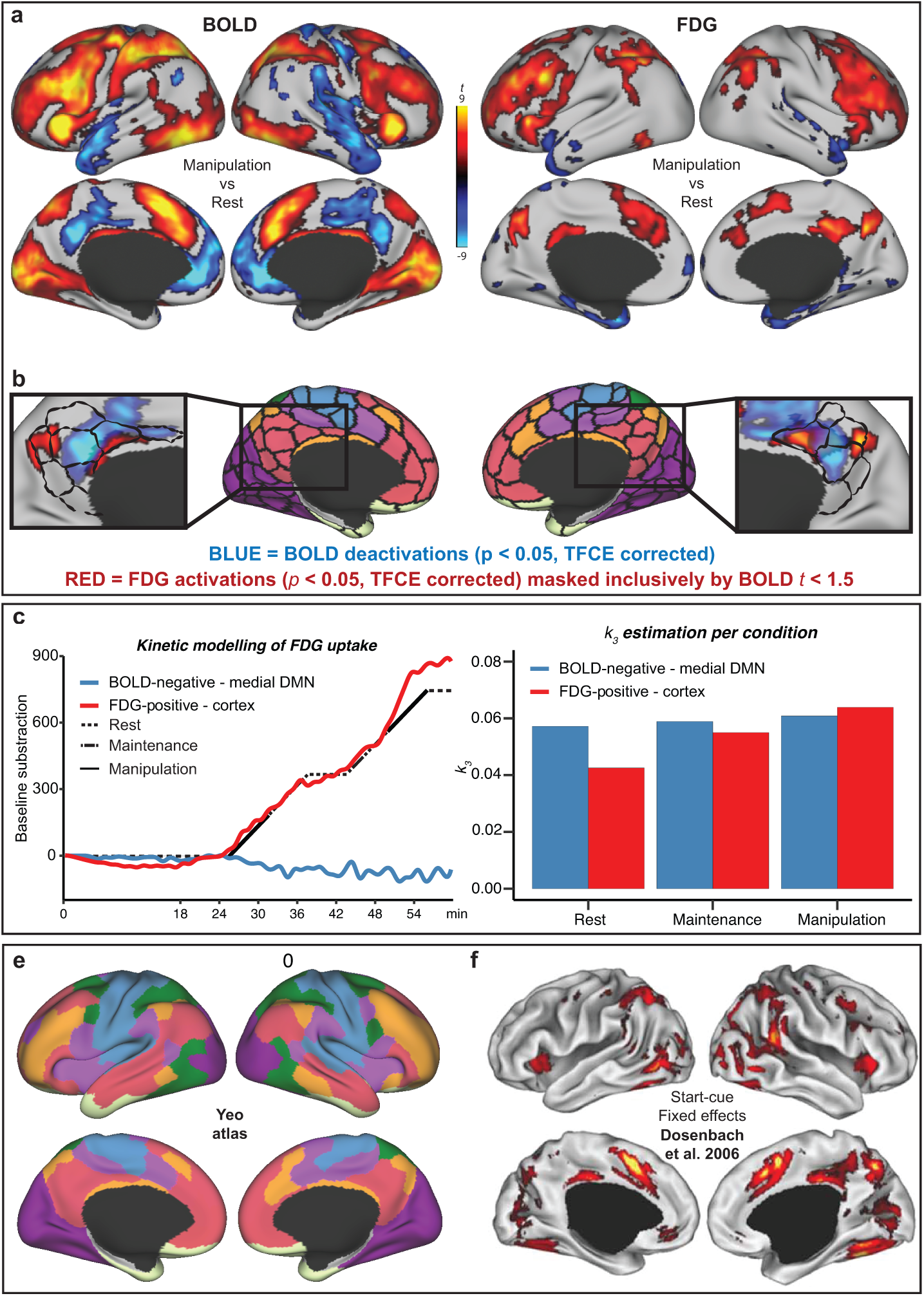
Divergence between the BOLD response and glucose metabolism in the posterior DMN. **a** *fMRI* and *PET* data for manipulation > rest in red and manipulation < rest in blue (p < 0.05, corrected). **b** The posterior DMN from the Schaefer 300 parcellation are delineated on magnified views of the medial surfaces. Blue = fMRI deactivations, red = FDG activations (masked to only include voxels where the BOLD *t* < 1.5). Note that part of the FDG increase in the right DMN is partly overlapping with fMRI deactivation (see darker violet shading in panel b, or compare the two right medial views in panel a). **c** Kinetic modelling of the FDG signal was performed in two large *ROIs* reflecting FDG activations (red, manipulation > rest) (cf panel a), and DMN (blue, manipulation < rest in the medial frontal and posterior DMN. Residual FDG signal in the ROIs after subtracting the estimated baseline curve is represented by the red and blue lines. The black line illustrates the task structure. **d** The estimated *k3* for each condition and ROI are depicted with bars. **e** The Yeo atlas is displayed to provide a point of reference for panels a-b and f. **f** Results from a meta-analysis of start-cue effects capture several areas where FDG activations are seen without BOLD signal changes (adapted from^2^ with permission from Elsevier).

The absence of FDG deactivations in midline and parietal cortex was explored further with kinetic modeling of FDG data using standard PET compartmental modelling^46^ in both simulations (Supplementary Fig 3) and real data. Modeling of the observed data confirmed that FDG data in regions of DMN BOLD task deactivations showed no notable change in *k*_*3*_ throughout the experiment (Figure 3c-d). Because kinetic modeling is not dependent on an a priori defined GLM, this shows that a lack of FDG change in regions of core DMN is not merely reflective of an ill-fitting a priori model. Conversely, modeling of the activations showed that *k*_*3*_ was steepest during manipulation blocks, followed by maintenance, and lastly by rest (bars in Fig. 3d). As suggested by high k3 throughout rest and task in DMN (and mean uptake, cf Fig 1), we thus interpret the lack of task-related decreases in glucose metabolism in core DMN not as a relative absence of brain activity but as metabolic activity that *remains high* when individuals transition from rest to task.

Finally, and unexpectedly, increases in FDG signal during manipulation > rest were observed in a set of regions located largely adjacent to fMRI deactivations in DMN (with partial overlap in right posterior midline). To illustrate that significant FDG activations without spatially overlapping fMRI activations in these regions did not reflect sub-threshold fMRI activity, the statistical map for FDG data in Fig. 3b was masked to remove any voxels where the fMRI statistic in the corresponding voxel was above the liberal threshold of *t* = *1*.5 (*p* < 0.15 uncorrected). With the aid of the recent Schaefer 300 parcellation^47^, which builds upon the original Yeo parcellation^45^, this figure shows that task-related FDG increases and BOLD deactivations tend to occupy largely separate parcels within the DMN, with FDG activations being prominent at the border between DMN and FPN. Task-related increases in FDG signal without corresponding changes in BOLD were also observed in both left and right angular gyrus following the same pattern, but not in anterior DMN (i.e. MPFC) or temporal cortex.

In summary, comparisons of spatial overlap across modalities established three different patterns of modality convergence distributed throughout association cortex; (i) overlapping activation in both modalities in widespread task-positive areas, (ii) BOLD deactivations without corresponding change in FDG in core DMN regions, and (iii) increases in FDG at the border of task-positive areas and posterior DMN, largely without overlapping fMRI task-related activation or deactivation. Of note, the location of FDG increases without corresponding BOLD signal changes, but adjacent to DMN deactivations, were highly reminiscent of a set of regions previously described in terms of their involvement in processing task onset cues^48^ (Fig. 3f), suggesting that these regions are indeed important for cognitive control but are not captured in the fMRI GLM. This prediction was explored further in a set of model-free fMRI analyses described next.

### The precuneus exhibits a distinct BOLD signature when transitioning between rest and task contexts

Task-free functional connectivity analyses were computed for three different seed ROIs (Fig. 4a): Left ACG in the FPN (from hereon referred to as ACG/FPN↑↑ with the arrows denoting that both modalities showed a positive task response), left PCC in the posterior DMN displaying a negative BOLD response and no significant change in FDG signal (PCC/DMN↓⨯), and left PCu, located in the posterior DMN where there was no change in BOLD but increased FDG signal (PCu/DMN⨯↑).

**Fig. 4.**
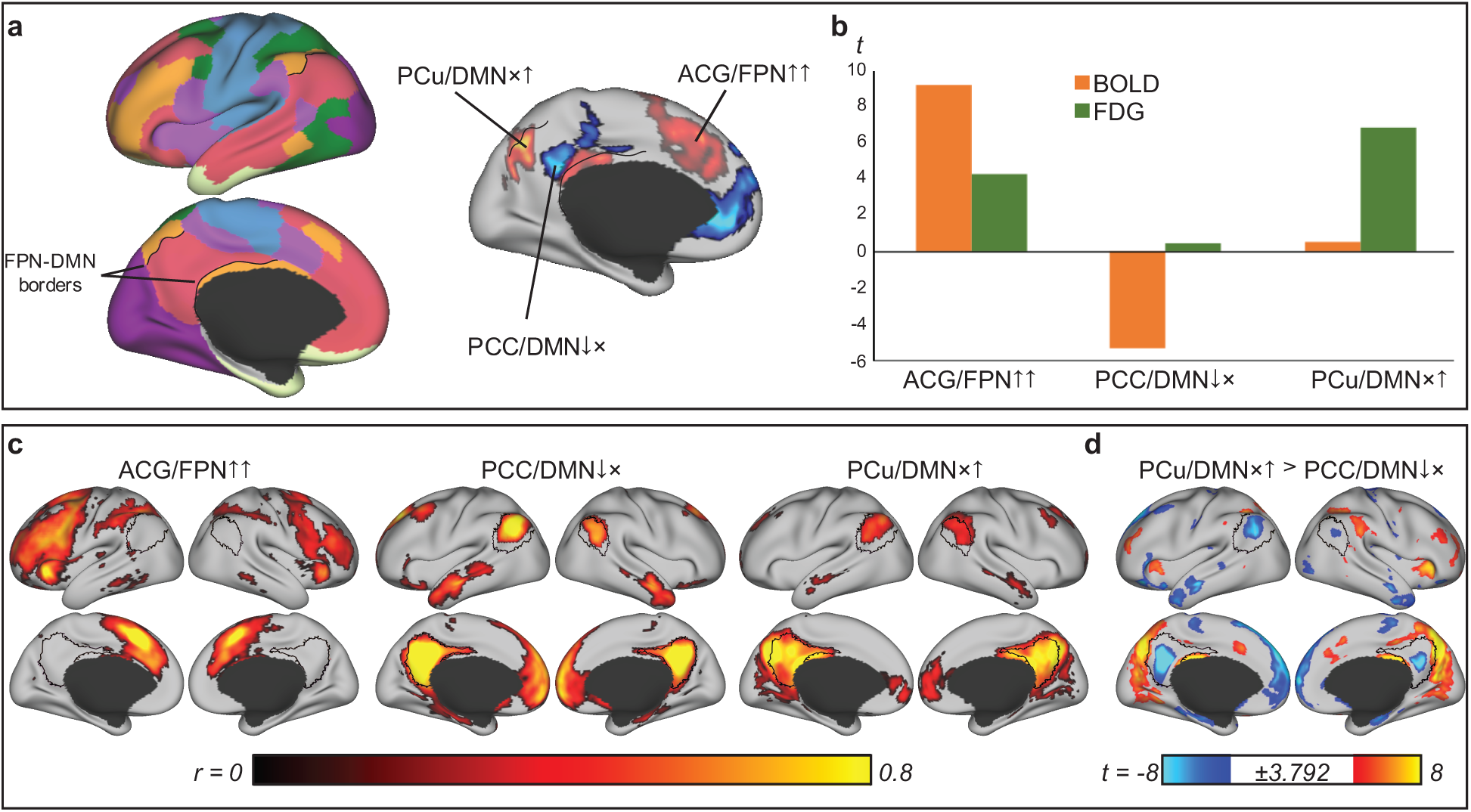
Functional connectivity during rest in three clusters exhibiting distinct multimodal patterns also display distinct connectivity patterns. **a** The Yeo parcellation is shown on the left with the borders between posterior DMN and FPN outlined in black. To the right, BOLD deactivations (manipulation < rest) are shown in blue, and metabolic increases (manipulation > rest) in red (opaque). Spheres with a radius of 5mm were placed in left paracingulate, where both modalities showed a positive response (ACG/FPN↑↑), in left posterior cingulate cortex where there was a negative BOLD response only (PCC/DMN↓⨯), and in right precuneus where there was increased metabolism only (PCu/DMN⨯↑). **b** The mean *t*-statistic within each cluster are depicted with bars for fMRI (orange) and PET (green). **c** Functional connectivity maps, *r*, from each seed to every other cortical voxel averaged across the three rest blocks. The functional connectivity maps were masked with maps derived from separate one sample *t*-tests performed on *r-to-z*-transformed images, thresholded at *p* < 0.001 (*t* > 3.792, two-tailed). **d** The contrast PCu/DMN⨯↑ > PCC/DMN↓⨯ reveal positive connectivity patterns along posterior DMN borders (thresholded at *p* < 0.001) and additional regions within attention networks also showing a start-cue effect in Fig. 3f. FPN = frontoparietal network, DMN = default-mode network

ACG/FPN↑↑ and PCC/DMN↓⨯ displayed connectivity patterns typical for their predicted network belonging, i.e. ACG/FPN↑↑ was coupled to regions of the FPN and DAN and the PCC/DMN↓⨯ coupled to regions of the DMN. In line with our description above of PCu/DMN⨯↑ being situated at the border of DMN, PCu/DMN⨯↑, connectivity in posterior midline extended beyond the DMN border to include cingulate cortex regions of the FPN as well as the border region between DMN and FPN in lateral parietal cortex (specifically, note that the positive connectivity is apparent also to the more distal PCC part of FPN and not only the neighboring FPN, delineated with black borders separating the posterior FPN and DMN networks in Fig. 4a,c). When contrasting PCu/DMN⨯↑ > PCC/DMN↓⨯ a set of regions emerges, including not only posterior DMN borders, but also additional frontal and parietal regions, and which again show striking resemblance to the set of regions meta-analytically found responsive to task start-cues^48^ (Fig. 3f). This shows that the three sets of regions identified by modality (non-)convergence described above each reflect intrinsically coupled networks, indicating functional heterogeneity also in the absence of task.

In a final set of analyses, we explore the BOLD signature that characterizes the PCu/DMN⨯↑ region in comparison to those regions where we observed a significant task-related change in the fMRI GLM analysis. The BOLD time-series of the ROIs were extracted and plotted for different task windows (Fig. 5). As is evident from a visual comparison of BOLD signatures, PCu/DMN⨯↑ is characterized by sharp increases at the onset of each new block. These are particularly pronounced at the first task onset, i.e. after switching from a long rest period to task (find additional violin plots and associated statistical tests in Supplementary Fig. 4). The time-series of correlations and lagged correlations in Fig. 5d across this window suggest that PCu/DMN⨯↑ is functionally correlated to ACG/FPN↑↑ around transitions from rest/fixation to task and early into tasks, whereas it is highly correlated to PCC/DMN↓⨯ towards the end of the trial period. Intriguingly, higher lagged correlations indicate that changes in PCu/DMN⨯↑ *precede* signal changes in ACG/FPN↑↑ and PCC/DMN↓⨯ both at the beginning and at the end of the task. This lag was not due to the application of bandpass filtering as the lag remained also for non-filtered BOLD time-series (Supplementary Fig. 5). From these data, we conclude that the reason that the metabolic activity in PCu region was not paralleled by an activation in the fMRI GLM is that the underlying BOLD time-series in these border regions did not correspond to a boxcar model of task onset and offset. Instead, it preceded FPN activations at the beginning of the task block, and then switched to preceding DMN deactivations during the latter half of a task block. This also shows that activity in regions situated at the border of FPN and posterior DMN does not reflect a blurred signal of two networks but rather flexible coupling to one or the other network during the task window, tentatively reflecting a continuous “gating” in neighboring networks. Supplementary Fig. 6 reports additional fMRI time-series analysis in striatum where FDG activations without significant fMRI activations were observed in medial caudate, areas associated most strongly with DMN^49^. The pattern clearly resembles that observed in posterior DMN in cortex, implicating also striatal regions in guiding BOLD signal increases and decreases during cognitive control.

**Fig. 5.**
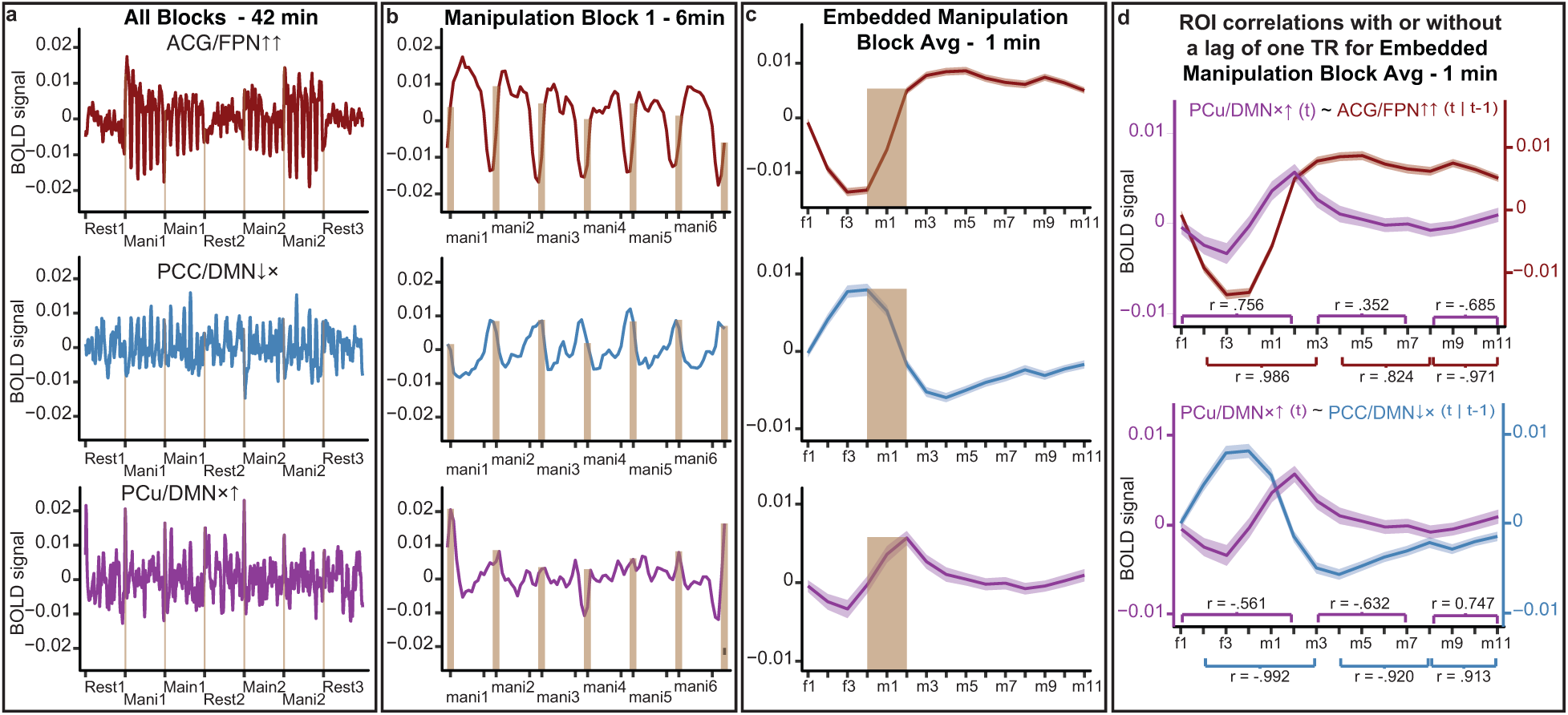
The BOLD time-series in the metabolically active posterior precuneus displays large amplitude shifts at transitions between blocks and precedes BOLD signal changes in ACG/FPN and adjacent PCC/DMN. **a**-**c**. The BOLD time-series for the three cortical ROIs exhibiting distinct multimodal signatures, were averaged across participants. TRs with a framewise displacement above 0.25 mm were removed from analysis. The block types in **a-b** were centered on the midline of each block/event, whereas the event TRs in **c-d** were centered on the onset. To aid visualization, brown bars were placed on the transition zones between blocks in **a**, and when transitioning from fixation to task periods in **b**-**c** (the one TR prior to the transition and the two subsequent TRs). The height of the bar was set based on the local minima/maxima observed along the bar’s width. **a** The BOLD time-series plotted across the 42 min fMRI paradigm. **b** The BOLD time-series is shown for the first manipulation block, consisting of six shorter embedded blocks with 45 s of task (mani1-6) separated by 15 s of rest periods (+). In **c**, the time-series of the twelve 1 min embedded manipulation blocks were averaged across each of the four resting fixation TRs [f1-f4] preceding each embedded task block, and the eleven TRs [m1-m11] involving manipulation trials. **d** The averaged time-series from PCu/DMN⨯↑ in **c** correlated with the time-series in ACG/FPN↑↑ (top) and PCC/DMN↓⨯ (bottom) without lag (*r* presented above the x-axes) or shifting the two latter ROIs one TR backward (*r* presented below the x-axes). The time-series were split into a transition phase [f1-m2], a middle phase [m3-m7] and a late phase [m8-m11]. For the late phase, the lag consisted in shifting PCu/DMN⨯↑ to [m7-m10] and comparing those TRs to m8-m11. Note: the time-series without band-pass filtering shifts the transition related change approximately one TR forward, but the lag remains and is not simply an artefact from filtering. ACG = *anterior cingulate gyri*, DMN = *default-mode network*, FPN = *frontoparietal network*, PCC = *posterior cingulate cortex, PCu* = precuneus, *TR = repetition time*

## Discussion

In this study, we used simultaneous fPET-fMRI with a continuous infusion of FDG during alternating blocks of rest and WM to assess whether changes in blood oxygenation in response to a controlled task are paralleled by changes in glucose metabolism. With FDG being a proxy for synaptic activity that is independent of blood flow, we reasoned that the results inform us whether fMRI deactivations during an externally focused cognitive control task are reflective of a decrease in synaptic activity in the DMN. Our first major conclusion from these data is that this is not the case; unlike the BOLD signal, FDG signal in the DMN was as high during rest as it was during task. The disconcordance between blood oxygenation and glucose metabolism was specific to DMN as the two signals displayed increases during task versus rest in a canonical WM network including fronto-parietal regions involved in cognitive control and attention. The BOLD and FDG signals were also similarly modulated by task difficulty in these areas, with greater signal changes for a more demanding WM condition, and deactivations in the temporal cortex of DMN partly aligned between modalities, suggesting that the absence of deactivations in DMN are not due to an unknown technical limitation. These results are not compatible with the idea that DMN BOLD deactivations during task reflect a role for the DMN that is restricted to internal modes of cognition that are dominant during rest (e.g. autobiographical thought and mind wandering).

A second major discovery in the current study was that of task-related increases in FDG signal in regions located in posterior DMN, adjacent to BOLD deactivations. Analyses of fMRI time-series data in these regions suggested that these border regions guide activity in the neighboring networks during task, possibly signaling the switch between rest and task contexts. Below, we draw parallels between our results and invasive recording studies to provide putative mechanisms for our results.

### Comparisons with electroencephalography

The limited number of animal studies on DMN deactivations have failed to show clear evidence that BOLD task deactivations in DMN are reflective of relative decreases in neural activity during externally focused cognitive control^31–35^. For example, despite reduced *local field potentials* (LFP) across frequency bands in the cat PCC during a sensory discrimination task, an increase in neural firing was observed^31^, counterintuitively suggesting that neural activity increases may result in task-negative responses during externally guided attention. Similarly, while there is evidence for suppression of gamma band activity in the macaque analogue of PCC, concomitant increases in lower band frequencies have been observed during an attention task^34^. In line, intracerebral electroencephalography (iEEG) recordings in patients during surgery found that DMN regions showed reductions primarily in gamma band activity^53,54^, however, only a subset of electrodes placed within the DMN exhibited an antagonistic response when comparing rest or internally directed tasks to externally focused cognitive control tasks. These data suggest that while BOLD deactivations may be consistent with decreases in gamma band activity, changes in gamma band activity alone do not provide a complete picture of DMN neural activity during externally focused cognitive control. The current findings of continuously high glucose metabolism during both task and rest support this view.

Gamma-band activity is the oscillatory pattern of neural activity that is most strongly and consistently associated with task-specific stimulus onset and increased vascular responses^26,50,52^. In contrast, negative associations between oscillations and vascular responses have been observed in the middle frequency bands alpha and/or beta^33,50^. Moreover, gamma-band oscillations are associated with local processing, whereas the slower frequency bands are better suited for long-distance communication, e.g. from neuromodulation^55^. In this way, task-negative BOLD responses during cognitive control may be reflective of changes in gamma band activity associated with local processing, but BOLD signal changes likely disregard task-dependent changes in lower frequency bands. Indeed, using MEG^56^, DMN interactions to other networks were found strongest in beta, followed by alpha, which could serve to activate or suppress distal regions^50,55,56^. In accordance, stimulation using depth electrodes in resting human subjects revealed a stronger effect on FPN and salience network activity after DMN stimulation than vice versa^57^, a putative directionality also observed using granger causality or lagged correlations in rsfMRI data^58,59^. These observations are consistent with a more general role for the DMN in monitoring of internal and external environments alike^3,17,60,61^, or in the ongoing regulation and reconfiguration of brain dynamics and network states^4,15,16^. The posterior DMN in particular is a functional hub, with massive structural connections to other parts of the brain and connectivity analyses have demonstrated that the DMN is the most interconnected network in the brain using both fMRI^62^ and MEG^56^. This positions the posterior DMN in an ideal position to process and integrate transmodal information, also unrelated to specific sensory input.

In sum, the present finding that the DMN is metabolically active during both task and rest is consistent with the interpretation that the DMN is not one network but rather a heterogeneous set of regions^15,62–65^, with BOLD deactivation being reflective of only one facet of this heterogeneity. More specifically, within the DMN, activity might shift from regions involved in stimulus-specific, or local processing to regions involved in context-specific processing, or distal network communication during cognitive control, with task-dependent changes in BOLD signal preferentially sensitive to the former. This interpretation does not preclude the detection of differences in glucose metabolism in DMN in different task settings. For example, in a comparison between rest and a task specifically taxing internal modes of cognition in DMN (e.g. autobiographical memory), increases in FDG should accompany increases in BOLD. Similarly, prior work using simple sensory tasks without high cognitive demands, e.g. visual stimulation or finger tapping^40,42^, has suggested decreases in glucose metabolism in DMN areas.

### Increased glucose metabolism during WM adjacent to BOLD deactivation

Activity related to a shift in context has been observed previously for regions belonging to the DMN in both humans^17,48^ and macaques^32,33^. BOLD data collected from 8 separate studies involving tasks of decision-making, reading and verb generation, were used to identify a set of regions responding specifically to start cues in at least 80% of the tasks^48^. This set of regions (Fig. 3f), shows striking similarities to the network of regions along the borders of posterior DMN identified as being metabolically active during task in the current study, i.e. PCu, PCC, lateral parietal, and angular gyrus. In the current study, we show that, (i) this set of regions formed an intrinsically connected network, situated directly adjacent to BOLD deactivations in posterior DMN, (ii) glucose metabolism in these regions was high throughout the entire task block and not restricted to task onset, suggesting ongoing activity during task, and (iii) while the beginning of a task block was indeed characterized by a sharp increase in BOLD activity, the time course in later task periods was not random but coupled to the DMN regions that show a BOLD decrease during task (Fig. 5). Intriguingly, (iv) activity in this network preceded the BOLD increase in FPN and decrease in DMN. Taken together, the current results extend a role for this network from processing task cues^48^ to one that may involve continuous gating of activity in neighboring networks. As discussed for the disconcordance between FDG and BOLD signal in the preceding sections, the identification of heterogeneity of FDG responses to task within DMN supports the conclusion that there is high functional heterogeneity within the DMN^4,14,62–64,66^, that is not fully captured by task-related BOLD changes.

Notably, we observed largely identical results in the striatal counterpart to DMN, displaying an increase in metabolism without a concomitant BOLD-signal increase, exhibiting a transient BOLD-signal increase preceding signal changes in adjacent associative and attentional parcels of striatum. The striatum is a subcortical hub interconnected with the cortex in “loops” that receive input from cortex and relay it back via thalamus^49,67,68^. A hypothesized role for the striatum in terms of a “gate keeper” in this loop, determining whether new information is allowed entry into cortical association areas, and guiding subsequent behavior^69^, is consistent with our interpretation for this network in continuous gating of activity in neighboring networks. In terms of large-scale neurocognitive architecture, and considering our discussion of task-related activity switches in DMN, continuous gating may relate to the reconfiguration of brain networks to achieve brain states adapted in accordance with context and ongoing goals^4^.

The interpretation of a continuous gating network that guides DMN is also compatible with known neurotransmitter mechanisms. The burst of fMRI signal at the beginning of the task in PCu/DMN⨯↑ may reflect activity in GABAergic neurons^70^ in order to reduce neuronal noise and quiesce adjacent networks that are not beneficial for the task at hand if excited^4,32^, i.e. local processing in DMN during cognitive control. A decrease in the GLU/GABA ratio may reduce spiking activity, whereas metabolically demanding synaptic activity involving ion pumping may still increase^71,72^, and may shift the network away from local processing and high frequency gamma activity, to slower beta band activity, potentially enabling nodes of the DMN to influence dynamics in distal targets^56^.

### Limitations

There are several limitations to this study. First, no arterial blood samples were collected, and absolute quantification of CMR_glu_ was therefore not possible. However, absolute quantification has been done to validate the GLM estimation of task-specific effects^40^. Thus, the pattern of task changes in the absence of CMR_glu_ quantification would not affect the conclusions drawn in the current study. Moreover, we did not manipulate activity in response to a task that has previously been shown to increase BOLD signal as compared to rest in DMN (i.e. autobiographical thought). Thus, the conclusions from the current study remain specific to the comparison between rest and WM and we cannot speak to metabolic changes in the DMN in response to other tasks that may further inform the cognitive operations rooted in the human DMN. Finally, the duration of task blocks may influence the appearance of metabolic decreases in DMN considering that decreases in anterior DMN were found only for 10 minutes, but not 1-5 minutes block durations^42^. Nevertheless, the current timing and task design elicited robust changes in FDG that paralleled BOLD changes in task-positive networks and there is no obvious reason why the design should conversely hinder the detection of activity in the reverse contrast.

### Conclusion

Using simultaneous FDG fPET and BOLD fMRI imaging during blocks of WM and rest we make several new discoveries that inform the interpretation of fMRI task deactivations in the DMN. Most importantly, the view of the DMN as a network with increased neural activity during passive states or internal modes of cognition only is not compatible with the current results. Unlike task-positive areas, the DMN is equally active in terms of energy consumption during both task and rest. We suggest that DMN activity during externally focused cognitive control may reflect a shift toward cross-network, long-range communication and orchestration across large scale association networks that BOLD is not sensitive to. These contextual shifts may, at least in parts, be driven by activity in a set of regions, most notably in posterior precuneus, that are situated along the border of the posterior DMN and the task-positive FPN. Going forward, future research should explore whether functional deficits in this putative continuous gating network mediate the blunted or absent task-induced BOLD deactivation observed in the DMN, in for example AD^21,23^, schizophrenia^25,73^, and aging^20,22^.

## Materials and Methods

### Participants

Participants were recruited via ads placed around the Umeå University campus and the city of Umeå and targeted healthy adults between 20-40 years of age with normal or corrected-to-normal vision. Exclusion criteria included history of head trauma, current or past diagnosis of a neurological or psychiatric illness, drug or alcohol abuse or dependence, and use of psychopharmaceuticals, drugs, or stimulants other than caffeine or nicotine for the past 6 months. Individuals having an MRI-incompatible metallic implant or object in their body were excluded for MRI safety reasons. Pregnant or breast-feeding individuals, as well as persons having previously undergone PET scanning for research purposes, were excluded for radiation safety reasons. Five participants were excluded due to technical problems in the acquisition or poor data quality, resulting in a final sample size of 23 healthy young adults (mean age = 25.2, SD = 4.0, range = 20-37, 56.5% female) that entered the analyses for the current study. This study was approved by the Regional Ethics Committee at Umeå University.

### Procedure

Participants were asked to fast for four hours prior to scanning. Upon arrival, participants were informed about the study, signed the informed consent form, and then practiced the in-scanner task. After practice, blood glucose levels were measured to confirm that levels were below 7 mmol/l, and an intravenous needle used for infusion was placed in the left arm. Infusion of FDG diluted in saline started at time zero of the PET-MR acquisition and continued for 60 minutes, distributing an initial radiation activity of approximately 180 MBq equally over the scan at a flow rate of 0.016ml/sec. MRI sequences, starting from time zero, were in order: MRI attenuation correction, T1-weighted structural, T2-weighted FLAIR, and, at the 18 minute mark, fMRI continuously for 42 minutes. At the 60 minutes mark, a B0 sequence was performed to acquire fieldmaps for fMRI bias field correction.

### Verbal working memory task

Verbal WM was assessed during fMRI using two conditions of a letter-based task, each performed twice during scanning in blocks of six minutes each (maintenance or manipulation of letters in WM). The order of task blocks was counterbalanced and interspersed with three six minutes rest blocks (Fig. 1A). The block order was rest, manipulation, maintenance, rest, maintenance, manipulation, rest. Six “fast” blocks that each contained five WM trials (45 s) and 15 s of resting fixation were embedded in each “slow” 6-minute task block. Whereas maintenance required only maintaining information in WM, the manipulation condition also required the active transformation of task information in WM. Robust fMRI responses to the task in canonical WM areas, FPN and DAN, have been reported in prior studies^74,75^. Prior to scanning, participants practiced four 1-minute blocks in each condition to ensure they fully understood the tasks. No participant performed more than two practice runs.

#### Maintenance and Manipulation

For the WM maintenance condition, each trial consisted of a target phase where four letters were presented for 2000 ms, a delay period of 3500 ms and a probe letter that was presented for 2500 ms during which participants were required to respond whether the probe corresponded to one of the target letters. Participants responded with their right index finger to indicate “yes” and with the middle finger to indicate “no”. Each trial was followed by a short fixation period of 1000 ms. In the manipulation condition, the timing was identical but the target phase included two letters the probe letter required participants to determine whether the probe corresponded to the subsequent letter in the alphabet of either of the two target letters. This condition requires maintenance and manipulation of the target in WM during the delay period.

### PET/MRI acquisition and analysis

All imaging was performed on a 3T General Electric Signa PET-MR system with a 16-channel head coil. PET coincidence data were collected and reconstructed to 60 1-minute frames using an OSEM (ordered-subsets maximization) algorithm with time-of-flight and point-spread function modelling, 2 iterations, 28 subsets, and 6.4mm post-filtering. Images were corrected for decay, scatter, and attenuation using an MR-based correction method (MRAC). The resulting voxel size was 0.97 ⨯ 0.97 ⨯ 2.81 mm^3^.

#### Positron Emission Tomography

The reconstructed FDG images were pre-processed in the following steps: (i) motion correction was performed by aligning each volume to the mean volume between minutes 40-45, (ii) spatial smoothing using an 8mm gaussian kernel, (iii) temporal smoothing using a running filter with a [0.5 1 0.5] kernel over three time points. (iv) normalization of tissue-activity curve (TAC) in each voxel to the maximum intensity of that voxel. After pre-processing, task-related changes in the slope of the TAC were analyzed using the *general linear model* (GLM) approach described by^39,40^.

Prior to task analysis, simulations of the task paradigm using a two-compartment pharmacokinetic model of FDG^46,76^, described in detail below, were performed to establish that the model would reveal kinetically plausible *k*_*3*_ changes in response to our task design and the task contrasts task > rest (i.e. activations) and task < rest (i.e. deactivations; Supplementary Fig. 3). In addition to establishing feasibility of the task design, this exercise indicated that a task-induced change in *k*_*3*_ would influence the measured PET signal by only a small fraction during the first two minutes because the increase in bound tracer is offset by a decrease of free tracer. In fact, a delay in the measurable change in FDG uptake in response to task onset can also be appreciated from the simulation figure in the original fPET publication^39^ but was never discussed in their work. Thus, separately for each condition, task blocks were modeled as a ramp function with slope = 1, delayed by 2 minutes into the task block. A baseline regressor was defined by a third-order polynomial across all gray matter voxels with the task regressors modelled as nuisance variables. The task regressors were then orthogonalized to the baseline regressor and added to the final GLM along with six nuisance movement regressors (x, y, z, jaw, pitch, roll). Finally, FDG task regressors were orthogonalized. However, considering that the manipulation task also requires maintenance of information in working memory, if added first into the GLM it may capture all common variance, despite orthogonalization. Accordingly, for the maintenance condition, we present statistics from a GLM with maintenance added first, and for the manipulation condition with manipulation added first.

The resulting beta maps were normalized to *Montreal Neurological Institute* (MNI) space using the T1 to MNI deformation field from the fMRI analysis pipeline. Higher-level analyses were performed using permutation testing with FSL randomise^77^ and 5000 permutations. A *threshold-free cluster-enhancement* (TFCE) threshold of *p* < 0.05 identified significant voxels. The main effects of manipulation or maintenance > rest as well as the contrasts manipulation > maintenance, and vice versa, were the contrasts of interest.

##### Kinetic modelling of dynamic FDG changes

While the GLM approach described above can identify relative changes in TAC slope as a function of condition, glucose metabolism cannot be quantified in an absolute sense without blood sampling. In order to address this potential shortcoming, we verified that the observed changes in task-induced FDG signal across association networks conformed to standard PET compartmental modelling (Fig. 3c) by quantifying k3 using the same FDG two-compartment pharmacokinetic model ^46,76^ that we used in the simulations. The arterial input curve from ^39^, which used the same dose of injected radiotracer as in the present study, was parametrized and used as the input-curve. The first 24 minutes post injection was a period of rest, which fulfills the assumptions for this model. A reference TAC (all gray matter voxels) from this rest-phase was fitted to the kinetic parameters K_1_, k_2_, k_3_, k_4_, fractional blood volume V_b_, and a blood-curve calibration-factor to accommodate the unknown scaling of the parametrized blood-input. It was found that V_b_ did not change the quality of the fit, so it was excluded.

The fitted parameters *K*_*1*_, *k*_*2*_, *k*_*3*_, *k*_*4*_ now define the rest-phase PET-uptake, *f(t)*. This curve, extrapolated to the whole scan duration represents the non-activated uptake. The amplitude of *f(t)* may in practice be scaled by region-specific differences in tissue accessibility and, maybe more prominently, differences in partial-volume effects. Describing such differences by a multiplicative factor, *s*, the measured PET uptake in the rest-phase can be described as *C*_*ROI,r*_= *s*. *f(t)*. The factor *s* also relates the ROI 0-24 minute area-under-the-curve for the rest-phase (*AUC*_*ROI,r*_) to that of the reference region (*AUC*_*ref,r*_), giving *AUC*_*ROI,r*_ = *s*. *AUC*_*ref,r*_. Eliminating *s* from these two relations yields

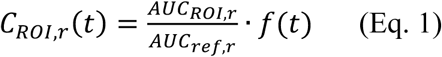

and represents the non-activated uptake-curve for the entire scan duration.

Baseline subtraction for a region was achieved by subtracting the time activity curve of the non-activated curve (Eq. 1) from that for the region, *C*_*ROI*_*(t)*

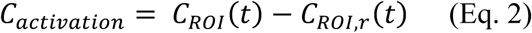

This alternative model approach was finally used in Fig. 3c-d to estimate the kinetics in task-positive and task-negative areas.

#### Magnetic Resonance Imaging

##### T1-weighted

Structural T1-weighted images were acquired for 7.36 minutes with the following acquisition parameters: [FOV: 25×20, matrix: 256×256, Slice Thickness: 1mm, Slices: 180, TE: 3.1ms, TR: 7200ms, Flip Angle: 12, Bandwidth: 244.1 Hz/Pixel]. T1 images were used for tissue segmentation and normalization to standard MNI space using a preliminary 12 degrees of freedom (DOF) registration with FMRIB’s Linear Image Registration Tool (FLIRT) followed by a nonlinear registration using FMRIB’s Nonlinear Image Registration Tool (FNIRT)^78^, resulting in 2 mm isotropic voxels.

##### fMRI

The BOLD data (Sequence parameters: FOV: 25.6, Matrix: 96×96, Slice Thickness: 3.6 mm, TE: 30 ms, TR: 4000 ms, Flip Angle: 80°, Acceleration Factor: 2.0) were pre-processed following conventional steps for fMRI as implemented in FSL FEAT (https://fsl.fmrib.ox.ac.uk/)^78^. Briefly, this included motion correction by volume-wise rigid body transformation to the first volume, slice timing correction, spatial smoothing (FWHM 5 mm), high-pass (120 s) temporal filtering and correction for B_0_ inhomogeneities using a study-specific fieldmap template based on 13 participants (data for 10 participants was lost due to an error in the acquisition). The magnitude and fieldmap volumes were normalized to standard space via T1 space and were then averaged to form study-specific templates. These templates were then back-transformed into each of the 23 subjects’ native space for use in FEAT. First-level statistics were obtained using the GLM. Manipulation and maintenance were modeled as separate regressors; regressors of no interest were the extended motion parameters, frame-wise displacement (FD), white matter signal, and cerebrospinal fluid signal. All resting periods (showing a fixation cross), including those within the embedded blocks, were treated as an implicit baseline.

Matched with the PET model, the main effects of manipulation or maintenance > rest as well as the contrasts manipulation > maintenance, and vice versa, were the contrasts of interest. The resulting volumes were normalized to standard space using the T1 to MNI deformation fields, and a 6 DOF boundary-based registration between the first EPI volume and T1. Higher-level analyses were performed with FSL’s randomise using 5000 permutations and treating a TFCE threshold of p < 0.05 as significant for a given voxel.

##### Regions-of-interest

To illustrate results and compute post-hoc analyses on fMRI time-series data, three ROIs, that each captured a distinct multimodal pattern observed in the GLM analyses, were defined along the left midline of the brain. Each ROI was produced by placing a sphere with a radius of 5 mm, i.e. five voxels in diameter, at three distinct locations along the midline: (i) In the left anterior (para)cingulate gyrus (ACG) located in the frontal FPN, both modalities displayed increased (↑) activation during performance of the manipulation task compared to either rest or maintenance conditions (ACG/FPN↑↑), MNI coordinates [−4 16 48]. (ii) In the left posterior cingulate cortex (PCC), located in posterior DMN, a decrease (↓) in BOLD was observed but with no (⨯) concomitant change in FDG signal (PCC/DMN↓⨯) [−6, −52, 28]. (iii) In the left precuneus (PCu), also located in posterior DMN, there was no change in BOLD according to the fMRI GLM but increased metabolism in the FDG GLM (PCu/DMN⨯↑) [−2, −72, 36]. For completion, a fourth ROI was defined in right PCC where there was a negative BOLD response and increased metabolism (PCC/DMN↓↑) [8, −44, 32]. The analyses of PCC/DMN↓↑ largely resemble that of PCC/DMN↓⨯ (data not shown).

In addition, three ROIs were also delineated in striatum by placing spheres with a radius of 3 mm, i.e. three voxels in diameter: (i) left dorsal pre-commissural caudate [−10 11 5] located in the striatal counterpart to FPN where both modalities increased activation during manipulation compared to baseline (dCau/FPN↑↑), consistent with cortical pattern of ACG/FPN↑↑, (ii) left anterior putamen [−26 7 −1] located in the *ventral attention network* (VAN) where an increased BOLD signal was observed without a significant concomitant increase in FDG (aPut/VAN↑⨯), and (iii) left anterior caudate [−10, 13, 1] located in the DMN where no BOLD signal change was observed despite increased metabolism (aCau/DMN⨯↑), consistent with cortical pattern of PCu/DMN⨯↑.

##### Functional Connectivity fMRI

The following additional pre-processing steps were performed for fMRI functional connectivity analyses: normalization of time-series, and application of a bandpass filter with the highpass set to 180 s and lowpass to 6 s, i.e. half the block length and two thirds the trial length). Each of the three 6-minute blocks of rest were separately processed. Pearson correlations were then performed between the average activity of each of the three cortical regions of interest (ROIs) to every other voxel in the brain. The resulting r maps were z transformed and then averaged across the three rest blocks. A one-sample t-test was then performed for each voxel and were considered significant at t > 3.792, p < 0.001. For visualization purposes, a binary mask of significant voxels was multiplied by the group averaged r map. A separate t-test was performed to contrast the connectivity maps of PCu/DMN⨯↑ and PCC/DMN↓⨯. The z map of PCC/DMN↓⨯ was subtracted from that of PCu/DMN⨯↑ for each individual subject before entered into analysis. The results were considered significant at −3.792 < t < 3.792, p < 0.001, two-tailed.

##### Time-series analysis

In order to visually inspect and explore the BOLD signal in the ROIs, the BOLD time-series, subjected to the same pre-processing steps as for the functional connectivity analyses, was extracted and averaged across participants. For illustration of condition-specific responses, the two “slow” manipulation blocks were averaged across participants, as were the twelve “fast” embedded manipulation blocks within these longer manipulation blocks, each consisting of four TRs of passively observing a fixation cross (f1-f4) and eleven TRs of performing the manipulation task (m1-m11). To reduce any influence of residual motion, TRs with a framewise displacement (FD) > 0.25 mm were excluded. In addition to visual inspection of the time-series, correlations were performed on transition periods between rest and task (f1-m2), midway through the task (m3-m7), and late into the task before the fixation period (m8-m11), each representing a different phase of the evolution of the BOLD response throughout the average embedded manipulation blocks. Correlations between time-series from different ROIs were performed using both moment-to-moment correlations and a lagged correlation with one TR delay imposed on ACG/FPN↑↑ and PCC/DMN↓⨯, where a higher correlation in the latter case would indicate that BOLD signal changes in PCu/DMN⨯↑ precede those in the other two ROIs.

## Acknowledgements

AR has received funding for this project from the European Research Council (ERC) under the European Union’s Horizon 2020 research and innovation programme (ERC-STG-716065). LR is recipient of a DOC Fellowship of the Austrian Academy of Sciences at the Department of Psychiatry and Psychotherapy, Medical University of Vienna. We thank the staff and leadership of Umeå Center for Functional Brain Imaging at Umeå University, Cancer Centrum and Nuclear Medicine at Umeå University Hospital for facilitating the data acquisition. For their contributions to data collection, analysis or comments and discussions of preliminary results we thank in particular Lars Nyberg, Helen Lindberg, Peter Young, Linda Douw, and Valentin Riedl.

