## Supplementary for "No metabolic “default mode” of human brain function"

### Supplementary Information

Supplementary Table 1. List of peak activation changes in fMRI and PET

Supplementary Fig. 1. Dice overlap between BOLD and fPET statistical maps

Supplementary Fig. 2. Reduced FDG uptake in temporal cortex during task

Supplementary Fig. 3. FDG simulations

Supplementary Fig. 4. Distribution of the BOLD data in ROIs, averaged across manipulation blocks

Supplementary Fig. 5. Effect of filtering on raw time-series inspection

Supplementary Fig 6. FDG and BOLD activations in striatum

#### Supplementary Data Table 1.

Peak activation foci for the manipulation > rest contrast.

| fMRI | | | | | | PET | | | | | |
|---|---|---|---|---|---|---|---|---|---|---|---|
| **Region** | **MNI coordinate** | | | ***t*** | ***k*** | **Region** | **MNI coordinate** | | | ***t*** | ***k*** |
| Cerebellum | 36 | -50 | -30 | 14.1 | 25615 | IFG[1] | 46 | 26 | 20 | 10.4 | 4218 |
| Intracalcarine cortex | 14 | -72 | 10 | 6.76 | | Precentral/MFG[1] | -38 | 6 | 30 | 8.4 | 3159 |
| Insula[1] | -32 | 20 | -2 | 12.7 | 18614 | SPL[1] | -28 | -54 | 40 | 7.3 | 833 |
| Precentral/MFG | -42 | 6 | 30 | 8.71 | | ACC[3] | -6 | 18 | 32 | 9.8 | 783 |
| Paracingulate | -4 | 14 | 46 | 7.75 | | Precuneus[4] | 4 | -68 | 34 | 8.7 | 620 |
| Paracingulate | 8 | 28 | 32 | 7.93 | | Lateral Occipital superior[1] | 38 | -68 | 48 | 6.8 | 612 |
| Putamen | -24 | 2 | 6 | 8.03 | | PCC[4] | 4 | -42 | 34 | 7.5 | 552 |
| Thalamus | -10 | -22 | 10 | 7.5 | | Insular cortex[3] | -30 | 24 | 4 | 7.9 | 522 |
| MFG | -32 | 0 | 60 | 7.35 | | SPL/Angular gyrus[2] | 32 | -48 | 42 | 6.5 | 352 |
| ACC | -8 | 24 | 28 | 6.58 | | Caudate[5] | -12 | 14 | 2 | 7.9 | 345 |
| Caudate | -14 | 0 | 18 | 6.76 | | Caudate[6] | 14 | 14 | 8 | 4.7 | 262 |
| Supramarginal gyrus | -32 | -46 | 40 | 8.53 | | ACC[1] | 6 | 30 | 28 | 7.8 | 173 |
| Lateral occipital superior[2] | 30 | -66 | 36 | 6.02 | 1360 | Temporooccipital cortex[2] | -50 | -60 | -16 | 8.3 | 114 |
| | | | | | | Lateral Occipital superior[2] | -26 | -66 | 40 | 5.6 | 87 |

All coordinates are reported in *Montreal Neurological Institute* (MNI) space. Peak locations were extracted from *t* statistic maps masked using a *threshold-free cluster-extent* (TFCE) correction of $p < 0.05$ and a gray matter probability map of 0.5. To achieve separation of the large fMRI clusters an additional threshold of $t > 5.402$, $p < 0.00001$, was set, and the first ten non-cerebellar clusters are reported below the original cluster. The anatomical naming convention was based on the Harvard-Oxford atlas. *k* = cluster extent, functional networks [1]-[6] defined by Yeo[1] if cortical and Choi[2] if striatal, [1]FPN, [2]DAN, [3]VAN, [4]DMN, [5]DMN striatum, [6]associative striatum, ACC = anterior cingulate cortex, fMRI = functional magnetic resonance imaging, MFG = middle frontal gyrus, PCC = posterior cingulate cortex, PET = positron emission tomography, SPL = superior parietal lobule.

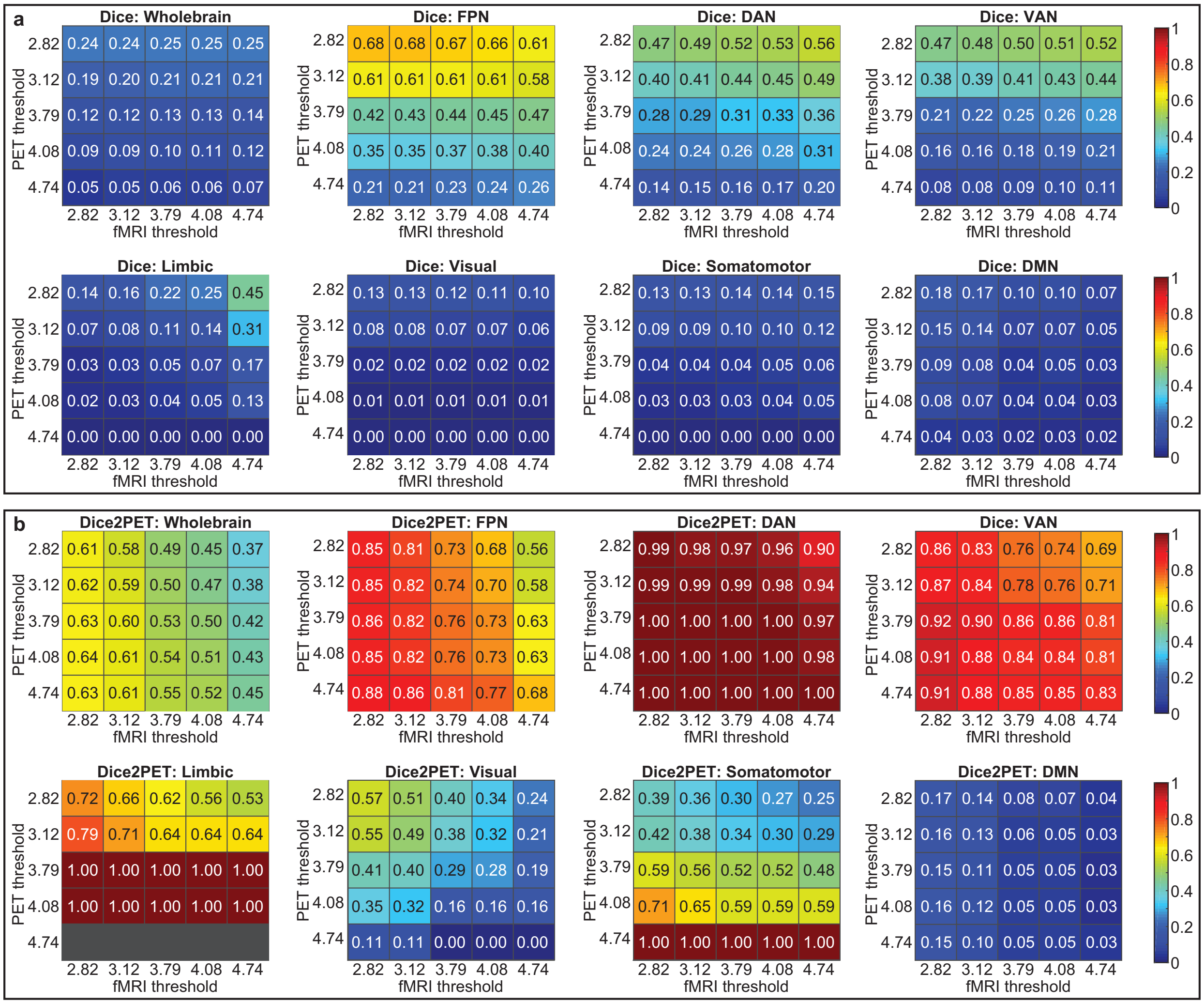


Supplementary Fig. 1. Dice overlap between BOLD and fPET statistical maps of the manipulation > rest contrast differs depending on functional network. The seven network parcellation from Yeo[1] was used to define functional networks, along with a whole-brain region encompassing all gray matter voxels. Dice overlap was calculated between the BOLD and fPET *t*-maps at different combinations of uncorrected thresholds, corresponding to $p < 0.01$ ($t < 2.818$) to $p < 0.0001$ (4.738) in increments of $p < 0.05$. **a** Original dice overlap scores are shown. As BOLD compared to FDG activations were generally more widespread an additional overlap score considering the union over the total amount of active PET voxels was calculated in **b** for each network. Comparing **a** and **b**, the drastically higher overlap scores primarily in DAN, VAN, limbic, visual, and somatomotor networks in **b** indicates that the voxels with increased FDG overlap with those showing increased BOLD, but that FDG is more focal, and BOLD more widespread. Note, however that this pattern may in part also arise if the modelling statistics for FDG is simply lower than for BOLD, irrespective of physiology. Regardless, the multimodal coactivation patterns clearly differ between functional networks, where attention networks show the highest similarity, followed by sensory and motor areas. FPN = frontoparietal network, DAN = dorsal attention network, VAN = ventral attention network, Limbic = limbic network, Visual = visual network, Somatomotor = somatomotor network, DMN = default mode network.

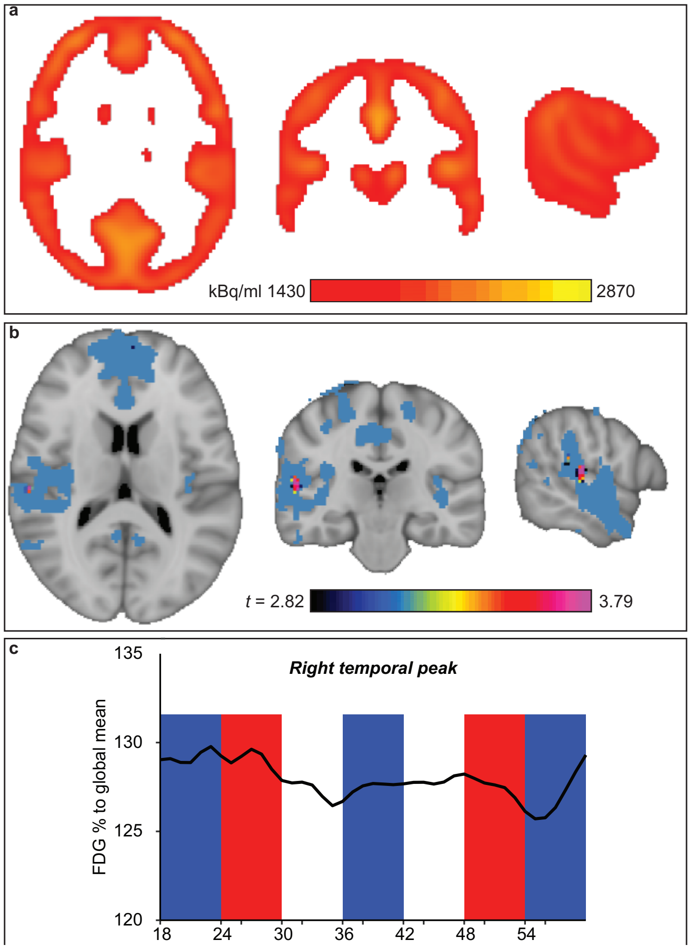


Supplementary Fig. 2. Decreased glucose metabolism during manipulation compared to rest in BOLD-task-negative regions was evident in the right temporal lobe. The whole-brain, corrected (tfce p > 0.05) voxelwise analysis revealed relative decrease in FDG uptake during manipulation compared to rest in bilateral temporal lobe. Because we did not have a-priori hypotheses about the specificity in temporal cortex and because it is an area prone to spurious result in both fMRI and PET, due to air/fluid-tissue boundaries and proximity to draining veins, we extended our analysis with a control analysis, restricted to voxels with high radioactivity uptake and showing robust BOLD deactivations. **a**. The FDG mask used for the analysis retained voxels with its maximum uptake during the full experiment above the $60^{th}$ percentile, essentially including only cortical grey matter not in the immediate vicinity of large draining veins or fluids [57 -25 12]. **b** The resulting *t*-test output was masked with significant BOLD deactivations ($p < 0.05$, *TFCE* corrected). With these additional precautions (which have no effect on the activations reported in the main text), FDG deactivations during task (manipulation < rest) were identified only in the temporal lobe, with a peak signal change significant at $p < 0.001$ ($t = 3.79$, uncorrected, two-tailed), and in anterior *medial prefrontal cortex* (MPFC). In order to be able to appreciate the peak and cluster-extent of the cluster, the threshold was lowered to $p < 0.01$ ($t = 2.82$, two-tailed) in this figure. The deactivation in MPFC was located at the edge of the mask however, running the risk of being an artefact and should be interpreted with caution. Conversely, the temporal cluster shown in **b** appears well within both the gray matter and BOLD deactivation masks. The fact that deactivations converge, at least at a commonly used uncorrected threshold, in temporal cortex shows that a lack of FDG deactivations in core DMN regions such as the posterior DMN is not reflecting a technical limitation to detect deactivations. **c**. Illustrated dynamic changes in FDG signal within the right temporal cortex peak [MNI coordinates 57 -25 12] showing increases in rest (blue) and decreases during manipulation (red). Because the kinetic modeling is sensitive to noise in small ROIs the timeseries illustration shows the % difference between FDG signal in the peak and the FDG signal in the rest of the FDG mask at each time frame, i.e. a model free illustration of dynamic changes in FDG distribution relative to the rest of the brain. A second tick mark was added 2 minutes after the tick mark illustrating the start of a new block.

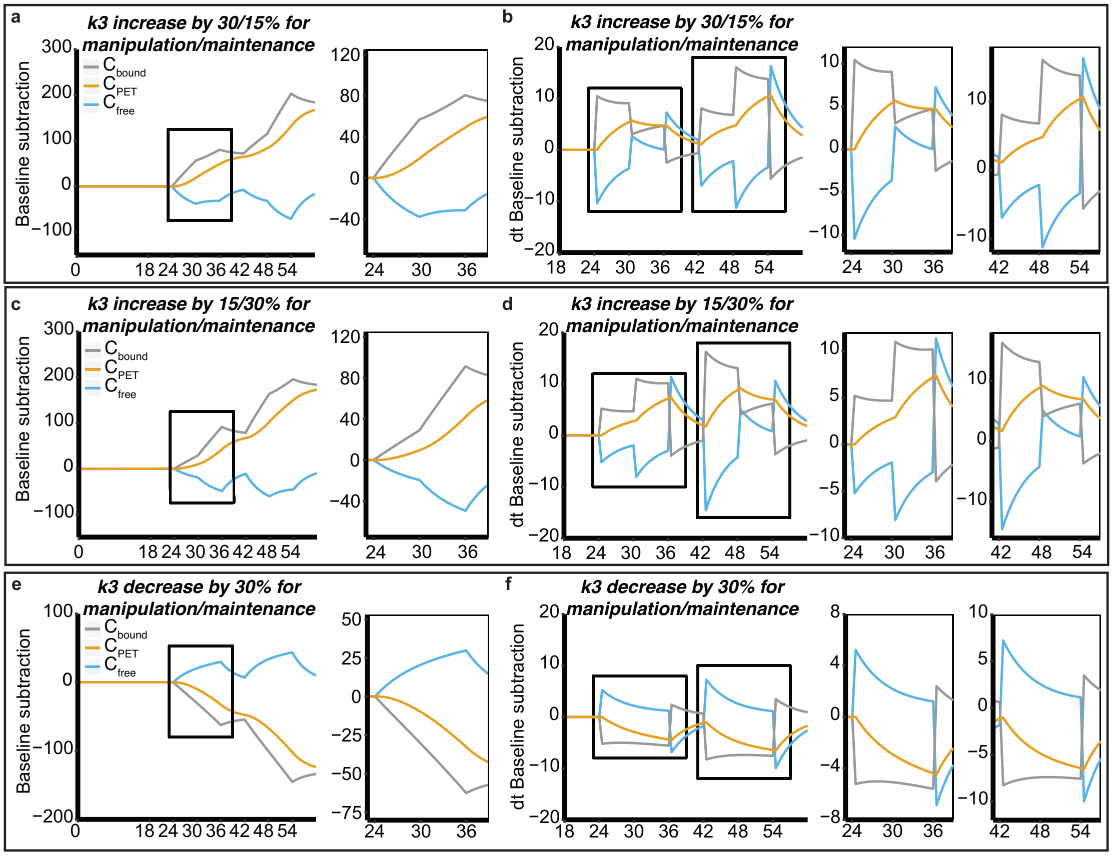


Supplementary Fig. 3. Simulations of metabolized, free, and tissue concentrations of FDG in response to blocked 6 min increases and decreases in $k_3$. The arterial input function and values for *K1* (0.1), *k2* (0.15), and *k3* (0.08) from[3] formed the basis for the simulations. $k_4$ was set to zero, and $V_b$ of 0.04 was used. To simulate the effect on the measured PET signal from differences in metabolic demand between conditions, $k_3$ was allowed to vary in blocks of 6 min. The concentration of tracer found in the tissue and blood ($C_{PET}$), corresponding to the signal measured in the PET scanner with real subjects, along with concentrations of free ($C_{free}$) and metabolized ($C_{bound}$) tracer was calculated for a 60 min long experiment (24 min of rest, followed by six 6 min blocks in the following order: manipulation, maintenance, rest, maintenance, manipulation, rest). The tissue concentrations under a baseline experiment (no change in $k_3$ during the 60 min) were subtracted from the concentrations in two activation and one deactivation experiment, resulting in a residual FDG signal reflecting increases or decreases in the rate of glucose metabolism under various conditions. **a** $k_3$ was set to increase by 30% under the manipulation, and 15% under the maintenance condition. An enlarged view of min 23 to 39 is provided as it illustrates the delay in $C_{PET}$ from a change in $k_3$ that is due to the interplay between tracer delivery, $C_{free}$, and $C_{bound}$. **b** The derivatives from **a** were plotted along with two enlarged views of both task periods as it clearly shows (i) that the change in slope for $C_{PET}$ in **a**, being the foundation for fPET, is not immediate (ii) that the slope increases for the full 6 min of a demanding block following a less demanding block, and (iii) that the slope remains positive even for subsequent rest blocks when k3 is equal to the baseline's. **c-d** the same analyses as in **a-b** were performed but with maintenance being the more demanding condition. In **d** it is apparent that the shape of the derivatives in the two panels are close to mirrored as compared to **b** with only minor differences. The scale is different however, suggesting that (i) the slope estimation in a given block differs slightly depending on the ordering of preceding blocks and (ii) that slope estimations will be higher in later blocks. Consequently, counterbalancing the order of blocks is important in order not to induce a systematic bias when comparing the magnitude of a change between conditions. **d-f** To simulate the response in areas where glucose demand is higher during rest compared to the tasks, $k_3$ was set to decrease by 30% during both the demanding and intermediate tasks. As in **a-d**, the sluggishness of $C_{PET}$ is evident, albeit in the opposite direction. Taken together these simulations show that, irrespective of the direction of $k_3$ change, the sluggishness of $C_{PET}$ should be taken into account when performing GLM based modelling of changes in the slope of $C_{PET}$. In theory, this could explain the reduced statistical sensitivity of fPET observed as blocks become shorter[4].

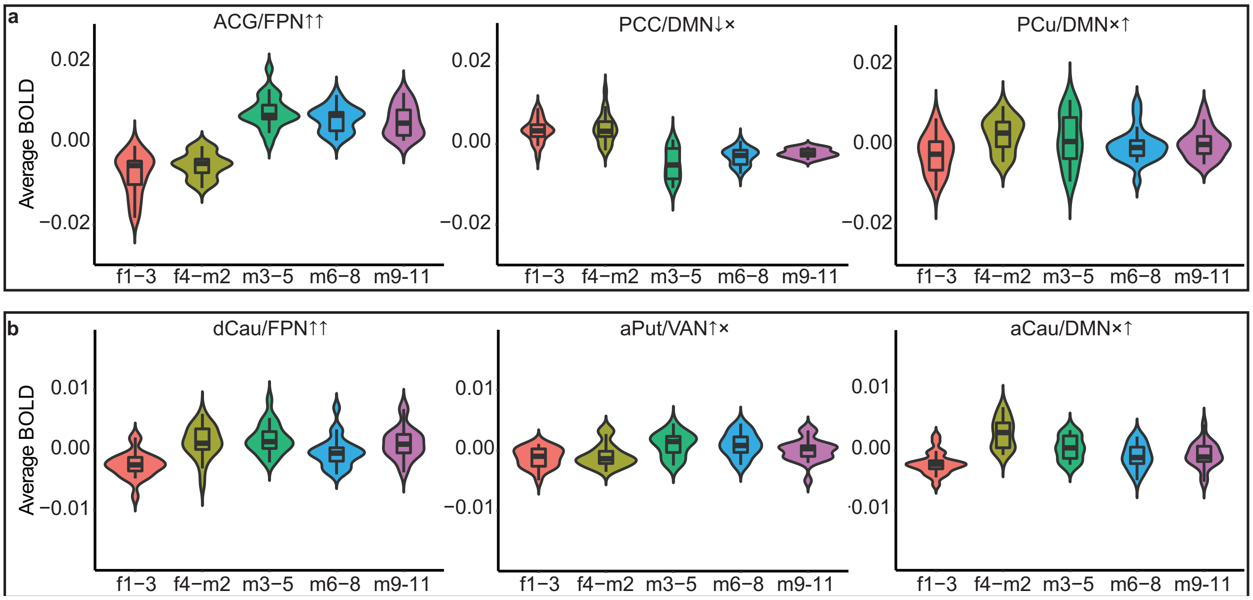


Supplementary Fig. 4. Distribution of the BOLD data averaged across manipulation blocks. For each participant, the BOLD signal in the twelve embedded manipulation blocks was averaged across every three TR in **a** the cortical ROIs from Fig. 4 and **b** the striatal ROIs from Supplementary Fig. 5. The violin depicts the density probability, and the box shows the median and interquartile range. When inspecting the time-series plots in Fig. 5 it was not immediately obvious why there was a significant BOLD activation (manipulation > rest) in dCau/FPN↑↑ but not in aCau/DMN×↑. When inspecting these violin plots it becomes obvious that the BOLD signal in dCau/FPN↑↑ remains above fixation levels throughout the 45 s of trials whereas the BOLD signal in aCau/DMN×↑ is strongly elevated a only after transitioning to task from rest before returning towards fixation levels. Paired *t*-tests comparing the transition phase (f4-m2) to the subsequent phases show no differences in dCau/FPN↑↑, whereas the transition phase is significantly higher for all comparisons in aCau/DMN×↑, all $p < 0.01$. aCau = anterior caudate, ACG = anterior cingulate gyri, aPut = anterior putamen, DMN = default-mode network, dCau = dorsal caudate, FPN = frontoparietal network, PCC = posterior cingulate cortex, *PCu* = precuneus, *TR* = repetition time*,* VAN = ventral attention network

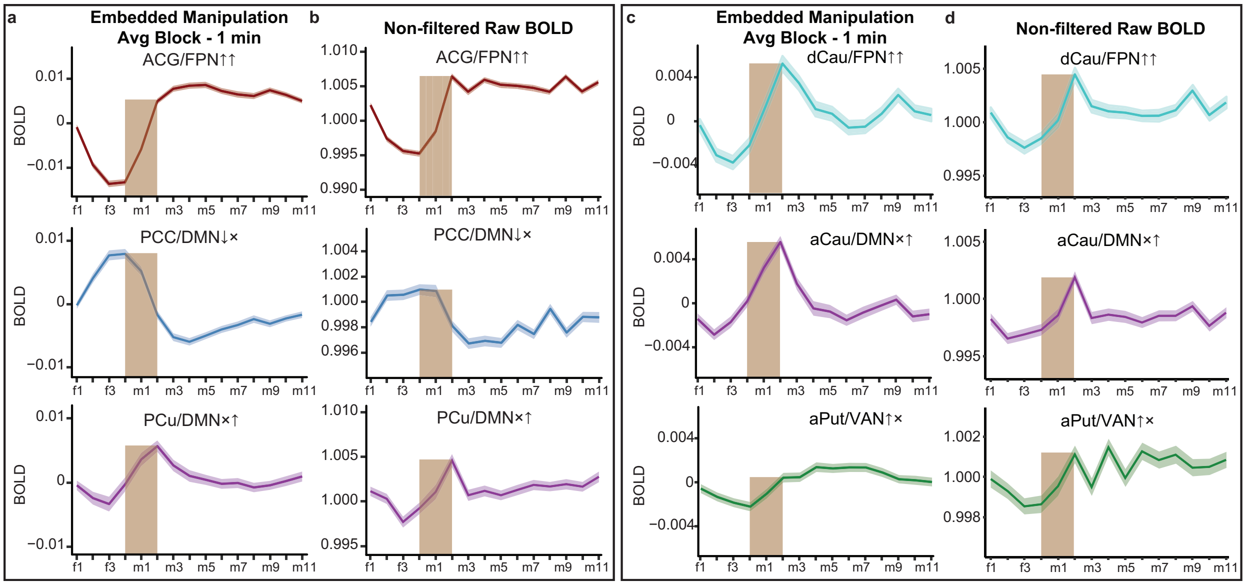


Supplementary Fig. 5. The filtered BOLD time-series from the ROIs in Fig. 4 and Supplementary Fig. 6 are displayed along with their non-filtered time-series. In **a,** the time-series of the cortical ROIs from Fig. 4 are shown. The twelve embedded manipulation blocks were averaged across the four resting fixation TRs (f1-f4) preceding each embedded block and the eleven TRs (m1-m11) involving manipulation trials. **b** The non-filtered BOLD signal is displayed, showing that the time-series lags are not artefacts from filtering. In **c,** the time-series of the striatal ROIs from Supplementary Fig. 6 are shown. **d** The non-filtered BOLD signal is displayed, showing that also the striatal time-series lags are not artefacts from filtering. aCau = *anterior caudate*, ACG = *anterior cingulate gyri*, aPut = *anterior putamen*, DMN = *default-mode network*, dCau = *dorsal caudate*, FPN = *frontoparietal network*, PCC = *posterior cingulate cortex*, *PCu* = precuneus, *TR* = *repetition time,* VAN = *ventral attention network*

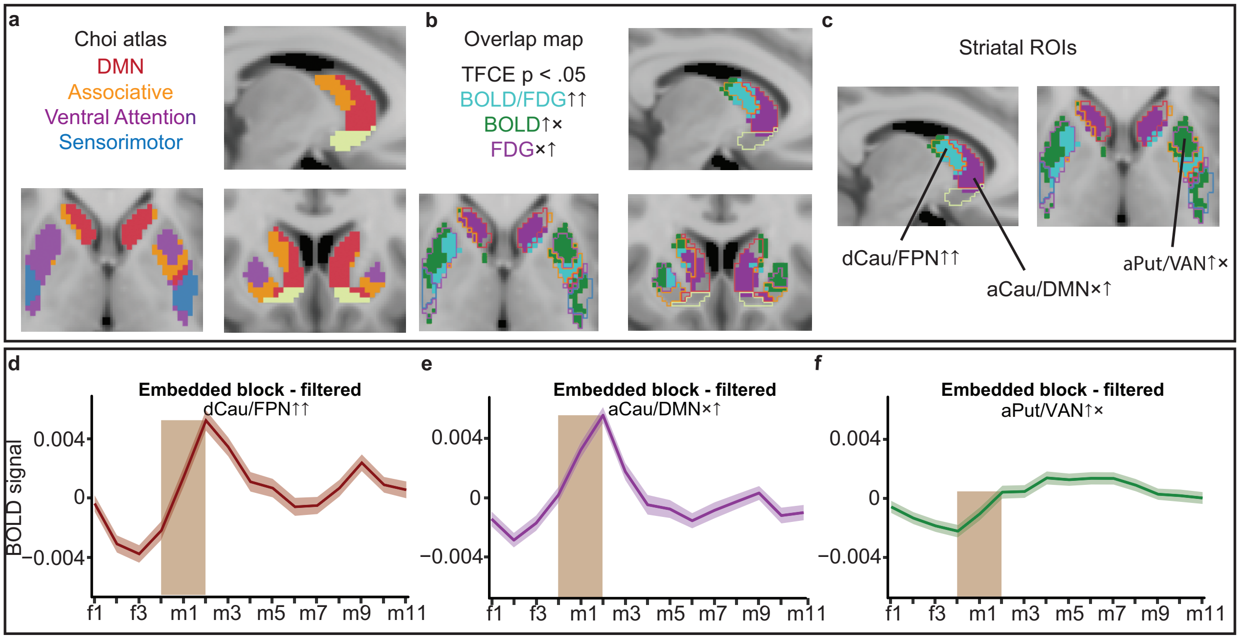


Supplementary Fig. 6. Differences between changes in BOLD and glucose metabolism in functional divisions of the striatum during working memory manipulation. The striatum is a structure with discrete functional areas. Choi and colleagues parcellated the striatum according to their belonging to the seven functional networks, including limbic, associative, attentional, and DMN parcels[5] (Supplementary Fig. 6a). This seven-network atlas was used for the subsequent comparisons between BOLD signal changes and glucose metabolism in striatum. Supplementary Fig. 6b below shows the overlap map between fMRI and FDG activations at $p < 0.05$ (corrected). Similar to the pattern observed in the cortex, BOLD activations and FDG activations overlapped in the dorsal striatum, associated most strongly with FPN. In sensorimotor putamen, BOLD activations were found but not significant FDG activations, likely reflecting the same pattern as generally observed in sensorimotor and motor cortex (i.e. sub-threshold FDG task activations). FDG activations without significant fMRI activations were additionally observed in medial caudate within areas associated most strongly with DMN, akin to the pattern seen in posterior DMN in cortex. However, in contrast to the cortical patterns presented in previous sections, no BOLD deactivations were observed within the striatal DMN. Inspection of the corresponding fMRI signal in the figure below shows, as in the cortical DMN, the BOLD signal change in medial caudate observed before the transition from rest to task precedes those observed in the other two clusters. The selected R-L views were positioned at MNI coordinates [-10, 14, 0]. **a** The Choi atlas[5]. The borders of the parcels are shown in **b-c** where BOLD and FDG activations and their overlap/non-overlap are presented. **b** Map of distinct multimodal BOLD and FDG signatures created by assigning colors to voxels based on co-activation or modality specific activations. **c** Location of the three striatal ROIs. (i) dorsal striatum where both modalities displayed increased activation, (ii) putamen where only BOLD increased, and (iii) medial caudate where only FDG increased. **d-f** The BOLD time-series during the twelve embedded manipulation blocks averaged across subjects (shaded areas represent ±1 standard error). To aid visualization, brown bars were placed on the transition zones between blocks when transitioning from fixation to task periods (the one TR prior to the transition and the two subsequent TRs). Of note, like in the cortex, at around the time of transitioning from resting fixation to task, the BOLD signal change in the DMN cluster exhibiting increased metabolism precedes the changes observed in the other two clusters. aCau = *anterior caudate*, aPut = *anterior putamen*, dCau = *dorsal caudate*, DMN = *default-mode network*, FPN = *frontoparietal network*, TFCE = *threshold-free cluster enhancement*, VAN = *ventral attention network*
